# Stoichiometric Insights into SARS-CoV-2 Spike–ACE2 Binding Across Variants

**DOI:** 10.1101/2025.04.04.647209

**Authors:** Ishola Abeeb Akinwumi, Sneha Bheemireddy, Laurent Chaloin, Serge Perez, Hamed Khakzad, Bernard Maigret, Yasaman Karami

## Abstract

The SARS-CoV-2 spike protein binds to the angiotensin-converting enzyme 2 (ACE2) receptor to mediate viral entry, with mutations in different variants influencing binding affinity and conformational dynamics. Using large-scale molecular dynamics simulations, we analyzed the Spike–ACE2 complex in the wild-type (WT), Beta, and Delta variants. Our findings reveal significant conformational rearrangements at the interface in Beta and Delta compared to WT, leading to distinct interaction networks and changes in complex stability. Binding free energy analysis further highlights variant-specific differences in ACE2 affinity, with alternative binding modes emerging over the simulation. The results enhance our understanding of spike–ACE2 stoichiometry across variants, providing implications for viral infectivity and therapeutic targeting.

## Introduction

Severe Acute Respiratory Syndrome Coronavirus-2 (SARS-CoV-2), is responsible for the highly contagious respiratory illness Coronavirus Disease 2019 (COVID-19) [1]. As of January 2025, this virus has continued to spread, resulting in more than 7 million deaths worldwide. Several variants of SARS-CoV-2, have been identified as Variants of Concern including Alpha (B.1.1.7), Beta (B.1.351), Gamma (P.1), Delta (B.1.617.2), and Omicron variants (B.1.1.529) [2]. Mutations in the SARS-CoV-2 genome, particularly in the spike protein’s Receptor Binding Domain (RBD) of the spike protein, contribute to variations in infection and transmissibility among these variants, as demonstrated in previous studies [3].

The surface of the SARS-CoV-2 virion is covered with spike protein trimers, primarily in the prefusion form, with three RBDs at the top. These trimers can adopt closed or open conformations, with the RBD occluded when in the down position. The spike protein contains a furin cleavage site, which modulates infection in a cell-type-dependent manner [4]. The subunits S1 and S2 of the spike protein play critical roles in recognizing and binding to host receptors and facilitating fusion between the host cell membrane and the viral envelope [5]. In fact, SARS-CoV-2 infection begins when the S1 subunit (RBD domain) binds to the ACE2 receptor on host cells. After receptor interaction, the Spike protein undergoes conformational rearrangement, exposing the S2 subunit, inserting the fusion peptide into the target cell membrane, and refolding into the postfusion form, resulting in viral membrane fusion [4]. The RBD is located in the closed state of the spike protein structure, making it unable to interact with the ACE2 receptor. This state is also the prefusion state, with S1 remaining atop the S2 subunit [6]. During the open state, the RBD interacts with the ACE2 receptor, initiating conformational changes that transition to the fusion state. The S2 subunit and the membrane control this reaction, leading to SARS-CoV-2 entry into cells and initiating the infection process [6]. Numerous studies have focused on the spike protein as potential therapeutic targets, including blocking receptor binding and developing DNA and RNA vaccines based on the S protein sequence [7].

While several studies investigated the conformational dynamics of the Spike-ACE2 complex and the variants of Spike, many questions remain to be answered. In this direction, molecular dynamics (MD) simulation has become invaluable for understanding the conformational heterogeneity and motion of macromolecular assemblies. MD simulations played a crucial role in vaccine development, RNA polymerase inhibitor design, RBD binding analysis, protease inhibitor design, and understanding the role of glycans in SARS-CoV-2 viral entry [8].

Here, we investigated the spike-ACE2 interaction networks in two variants (Beta, and Delta) compared to the wild-type (WT). For all studied systems, we considered a complex of Spike with three RBD up, and three copies of ACE2 embedded in the membrane [9] (**Fig. 1**), and performed 2-*µ*s long MD simulations to elucidate the stoichiometry between the two proteins over the simulation time. While our study is in agreement with the state-of-the-art on the set of key residue-players for the interaction between the two proteins, we highlighted significant conformational re-arrangements at the interface between the Delta and Beta variants compared to the WT complex, showing novel forms of interactions that strengthen the stability of the complex in these variants.

**Figure 1:**
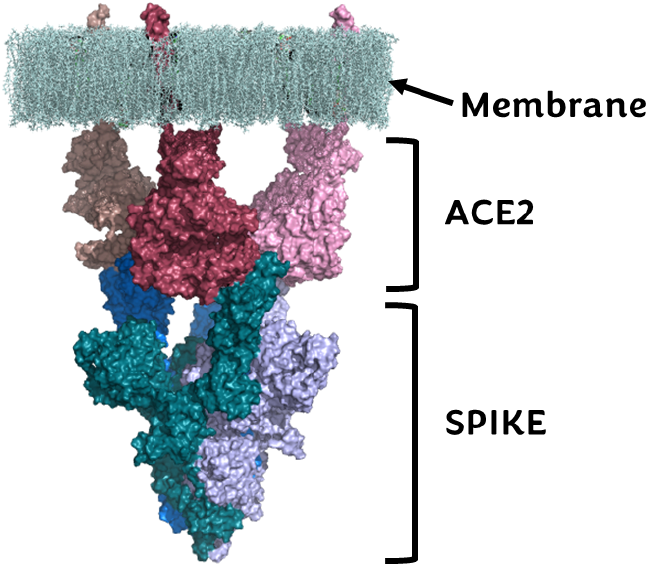
Spike Protein Trimer with three ACE2s. Spike trimer of WT complex with three ACE2 and membrane (Marine: chain A, Rhodium: chain B, Light Blue: chain C, Pink: chain D, Raspberry: chain E, Dark Salmon: chain F. chain A, B and C: Spike, while chain D, E and F: ACE2).

## Methods

### Preparation of Structures of Spike and ACE2

In this work we studied the conformational dynamics of the wild-type Spike trimer in a complex with three ACE2 proteins. For the WT (Wuhan-Hu-1 strain), the 3D coordinated were retrieved from the Cryo-EM structure with PDB code: 7KMS and the resolution of 3.64 Å. For the delta variant, the starting structure is considered the S-Delta variant (B.1.617.2) in complex with three ACE2-bound forms (PDB code 7V89, resolution 2.80 Å). Finally, for the Beta variant we started from the S-Beta variant (B.1.351) in complex with three ACE2-bound forms (PDB code: 7V7Z, resolution 3.10 Å).

### MD Simulations

Each experimental structure was embedded in a lipid POPC bilayer surrounding the ACE2 transmembrane tails and a water box. The box size of these systems was 300×300×400 Å^3^, and the total number of atoms was ∼4 million for each complex. In the case of the Delta variant, we also considered the glycans. For that we used the glycan-free model of the trimeric spike-ACE2 complex and built a new model of the Delta-ACE2 trimeric complex (Delta_*G*_) including now all the glycan groups for both spike Delta variant and ACE2 partners [10, 11]. After selecting the right glycans and their proper linkage to the proteins, we built the whole model using the CHARMM-GUI tool [12]. We used NAMD with the CHARMM36m force field for the MD simulations. Each model was first energy minimized with 64000 steps of conjugate gradient minimizer. Several rounds of refinement levels were obtained by first considering only the lipids and solvent as movable and the protein trimers as fixed, next relaxing the trimers’ side chains and finally releasing all constraints. Next, 10 ns equilibration steps were performed for each system before the 2 µs production phases. All simulations were carried out in the isobaric-isothermal ensemble, at constant temperature (300 K) and pressure (1 atm) using Langevin dynamics and Langevin piston as implemented in NAMD. The SHAKE algorithm was used to freeze bonds involving hydrogen atoms, allowing for an integration time step of 2.0 fs. For each studied system of WT, Beta, Delta and Delta_*G*_, we recorded 20,000 frames from the production trajectory (2 *µ*s; time step of 100 ps), leading to 8 *µ*s MD trajectories for further analysis.

### Analysis of the MD trajectories

#### Stability of the trajectories

We measured the root mean Square deviations (RMSD) over the C*α* atoms of each chain with respect to the initial frame of the production run. The RMSD of the Spike RBD and ACE2 without the trans-membrane region (ACE2_*noT M*_) were also measured. The results are shown in **Supplementary Fig. S1**, and average RMSD values are reported in **Supplementary Tables S1-2**. We also measured the per-residue root mean square fluctuations (RMSF) over the C*α* atoms and with respect to the average confirmation. The RMSF and MD simulations are reported in **Supplementary Fig. S2**, and average values are in **Supplementary Table S3-4**.

#### Pair interactions

For analyzing interactions between biomolecules, we have previously developed an approach referred to as pair interactions protocol [13, 14]. This is based on a feature of the NAMD package [15] allowing the user to change the standard output of a MD simulation so that the potential energy fluctuations between two non-bonded user-defined atom groups (referred to as a pair) are returned instead of the whole system’s energy. We used this analysis to quantify the interactions between the chains of spike and ACE2. Interaction energies, measured in kcal/mol, were calculated to assess the strength and stability of their binding.

#### Hydrogen bonds (H-bond) and salt bridges

H-bond analysis was performed to identify the interactions between the spike chains and ACE2 using the HBPlus program [16]. For each system, we reported H-bonds that were present at least in 40% of the frames along the simulation time. Salt bridges represent an essential class of non-covalent interactions, exerting a stronger force than a typical H-Bond due to the involvement of charged groups. These interactions occur between oppositely charged atoms, commonly found between the acidic and basic residues within proteins. In this study, the salt bridge interactions between the chains of the Spike and ACE2 for all the variants were analyzed. This analysis was performed using the VMD software [17]. We used a distance cutoff of 4.5 Å for the salt bridges calculation of all the variants and a percentage of 40% to get the salt bridges.

#### Binding Free Energy calculations

The AnalyseComplex in FoldX version 4 [18] was used to estimate the binding free energy between the chains of Spike and ACE2.

#### Clustering

To characterize representative conformations, we performed a cluster analysis of the MD trajectories using backbone RMSD as the similarity metrics by the GROMOS clustering approach [19] with a cutoff equal to 0.5 nm. Then the population of each cluster was calculated. Clusters with a population of at least 1000 conformations were reported.

#### Software

The gmx rms and gmx rmsf tool of Gromacs 2021.4 [20] software was used for the RMSD and RMSF calculations; CPPTRAJ in AMBER, was used to calculate the RMSD of the Spike protein’s RBD and ACE2_*noT M*_. We used the Saltbr package of VMD to obtain salt bridges. PyMOL was used to visualize the residues responsible for H-bonds and salt bridge formation.

## Results

### Hydrogen bond and salt bridge formation between the Spike and ACE2

To examine the spike protein’s interactions with ACE2, we analyzed and compared the H-bonds formed by all the variants. **Fig. 2A-C** illustrate the H-bond formations between chains A, B, and C of spike with chains F, E, and D of ACE2, respectively. We reported H-bonds present for at least 40% of the simulation times (see **Supplementary Table S5** for more details). The set of obtained H-bonds is mapped on the structure of each system (**Supplementary Fig. S3**) for a comprehensive overview of the H-bond network. We observed a specific interaction, K417-D30 in the WT between chains C and D (**Fig. 2C**), which was absent in the other variants. The Beta variant, intriguingly, exhibited H-bonds similar to those of WT, including G502-K353 (**Fig. 2A**), D405-K353 (**Fig. 2B**), and Q493-E35 (**Fig. 2B**) between chains A-F and B-E. Moreover, Beta had unique H-bonds, such as T500-Y41, R403-D38, and N487-Y83, which were not observed in WT. The Delta variant also shared H-bonds with WT, including G502-K353, D405-K353, Q493-K31, and Q493-E35. Notably, R403-D38 and Q493-K31 were present in chains B-E (**Fig. 2B**) and C-D (**Fig. 2C**) of the Delta variant, indicating their crucial role. Similarly, the Delta_*G*_ variant exhibited H-bonds Q493-K31, Q493-E35, G502-K353, and D405-K353, akin to those in WT. The G502-K353 H-bond was consistently present across all variants, underscoring its pivotal role in the spike protein’s interaction with ACE2. Differences between the Delta and Delta_*G*_ variants were observed in the H-bond formations. The Delta_*G*_ variant formed a unique H-bond, S477-M82 (**Fig. 2B**), absent in the Delta variant. Additionally, residue T500 in Delta formed H-bonds with Y41 and D355 of ACE2 (**Fig. 2A-B**).

**Figure 2:**
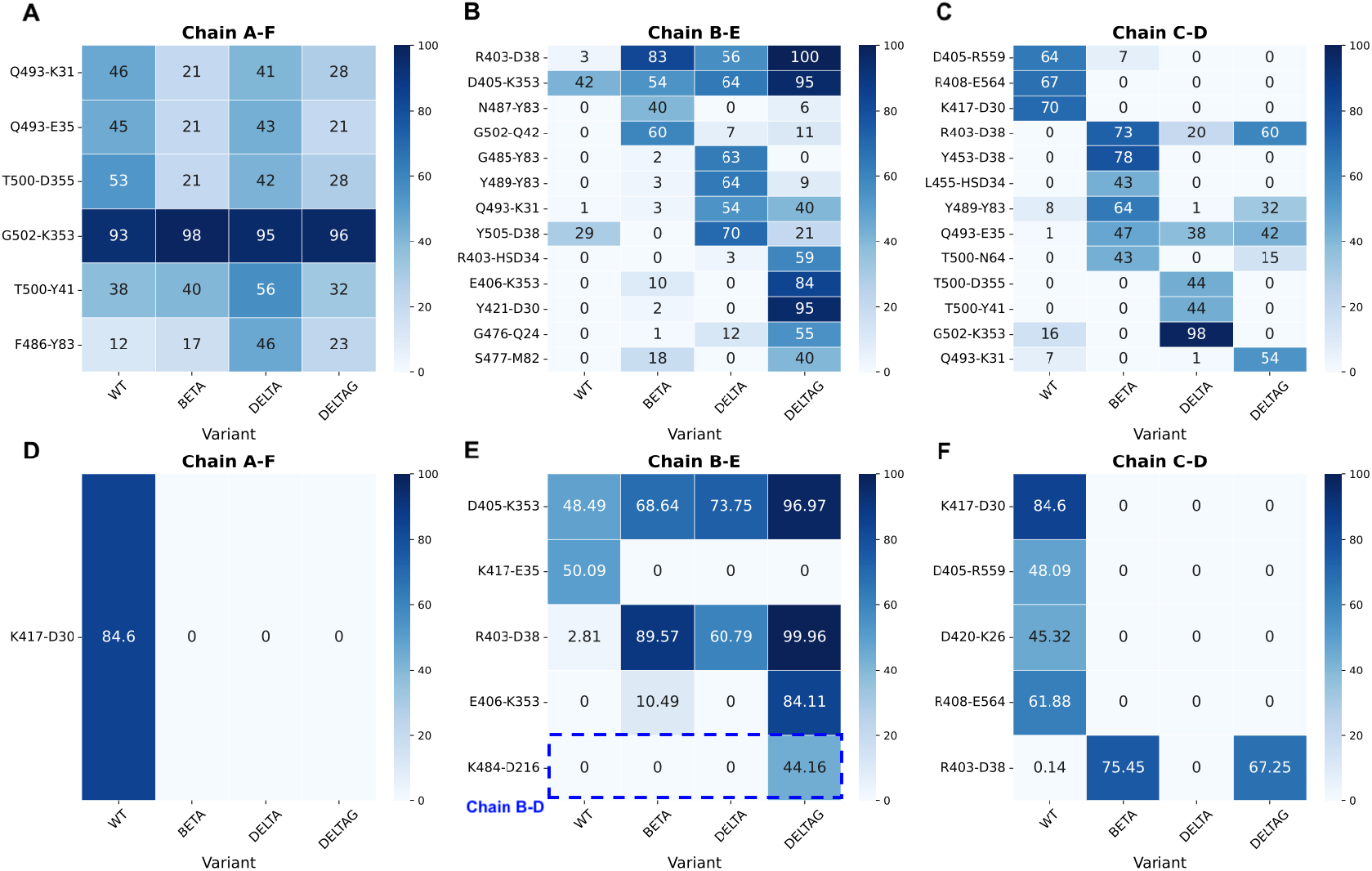
Key residues involved in the interaction network. H-bonds formed at the interface between the chains of spike trimer and ACE2 are reported for the WT, Beta, Delta and Delta_*G*_ variants (A, B and C). The stability of salt bridges at the interface is also reported for the WT, Beta, Delta and Delta_*G*_ variants (D, E and F).

We further analyzed the salt bridges formed between spike protein and ACE2. The details of the residues involved in those interactions are reported in **Supplementary Table S6**. Moreover, **Supplementary Fig. S4** illustrates the salt bridges mapped on the structure of the WT, Beta, Delta, and Delta_*G*_ variants. In the WT simulations, the K417-D30 salt bridge was only observed between chains A and F (**Fig. 2D**). No salt bridges were formed between chains A and F in the Beta, Delta, and Delta_*G*_ simulations. For salt bridge formation between chains B-E across all variants, D405-K353 was consistently observed. Delta_*G*_ exhibited a unique interaction between chains B-D (K484-D216) (**Fig. 2E**). The R403-D38 salt bridge was present between chains B-E in the Beta, Delta, and Delta_*G*_ variants and chains C-D in the Beta and Delta_*G*_ variants. Delta did not show salt bridge formation between chains C and D. These variations in salt bridge formation among the WT and other variants suggest structural changes that potentially impact the spike protein’s stability and its interaction with ACE2 receptors.

### Binding Free Energy

Evaluating the binding energies of the variants revealed differences in the interaction energy between the spike protein and ACE2. The average Δ*G* values are reported in **Supplementary Table S7** and color coded in **Fig. 3A**.

**Figure 3:**
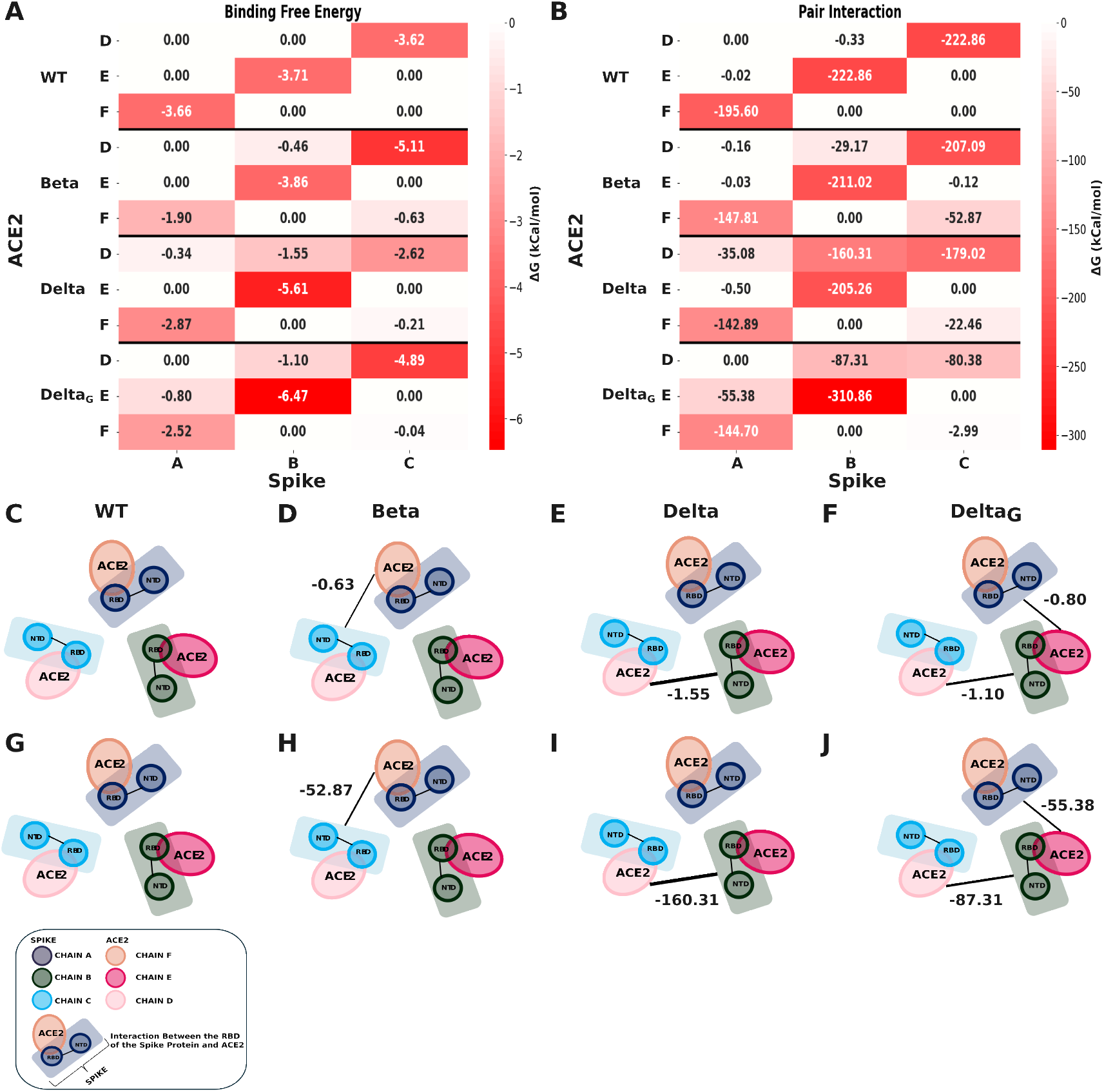
Binding free energy analysis. Binding Free Energy between the chains of spike trimer and ACE2 are reported for the WT, Beta, Delta and Delta_*G*_ variants (A), and Inter-protein interactions between the chains of spike trimer and ACE2 (C, D, E, and F). Pair Interaction between the chains of spike trimer and ACE2 are reported for the WT, Beta, Delta and Delta_*G*_ variants (B), and Inter-protein interactions between the chains of spike trimer and ACE2 (G, H, I, and J).

The interface between chain A of the spike and chain D and E of ACE2 (**Supplementary Fig. S5A-B**) showed Δ*G* values equal or close to zero for the WT, Beta, Delta and Delta_*G*_ variants, which could be explained by the fact that the corresponding chains are too far to have any binding affinity. Considering the interface between chain A of spike and chain F of ACE2, WT showed stronger binding affinity than the other variants (**Supplementary Fig. S5C**). Chain B of spike and chain D of ACE2 (**Supplementary Fig. S5D**) indicated lower Δ*G* values for the Delta, and Delta_*G*_ variants than WT and Beta variants. All variants exhibited favorable Δ*G* values between chain B of spike and chain E of ACE2 (**Supplementary Fig. S5E**), with Delta_*G*_ showing the strongest binding affinity with an average Δ*G* value of −6.47 *±* 1.94 kcal/mol (**Supplementary Table S5**). The interface between chains B and F (**Supplementary Fig. S5F**) showed zero energy values for all variants. The binding free energy between chain C of spike and chain D of ACE2 (**Supplementary Fig. S5G**) revealed that the Beta variant has a more substantial binding affinity, with an average Δ*G* value of −5.11 *±* 1.88 kcal/mol. In summary, all the variants represented stronger Δ*G* on average compared to the WT at the interface between the two proteins. While in the WT, only the direct interfaces (A-F, B-E and C-D) were involved, for the variants negative Δ*G* values were observed between neighboring pairs. Considering a cut-off value of −0.5 kcal/mol, we observed interactions between chains C-F in Beta, between B-D in Delta, and between A-E and B-D in Delta_*G*_. This is shown as a schematic representation in **Fig. 3C-F**, for all the studied systems. The black lines are drawn when the average Δ*G* is lower than −0.5 kcal/mol between a given pair of protein chains. This analysis is in agreement with the higher binding affinity between the spike and ACE2 in the variants compared to the WT.

With the pair interaction analysis, we measured the enthalpic energy between the chains of spike and ACE2, and the results along the simulation time are reported in **Supplementary Fig. S6**. The average pair interaction values are reported in **Supplementary Table S8** and shown as heatmap in **Fig. 3B** between the chains of spike and ACE2. The analysis between chain A of spike and chains D and E of ACE2 (**Supplementary Fig. S6A-B**) revealed no significant interaction for WT, Beta, Delta and Delta_*G*_. However, chain A in all the systems demonstrated interactions with chain F of ACE2 (**Supplementary Fig. S6C**). The pair interactions between chain B of the spike and chain D of ACE2 (**Supplementary Fig. S6D**) indicated interactions in the case of Delta and Delta_*G*_. All variants exhibited favorable interactions between chain B of spike and chain E of ACE2 **Supplementary Fig. S6E**), with Delta_*G*_ showing the lowest energy value of around −450 kcal/mol. We observed no interactions between the chain B of the spike and chain F of ACE2 (**Supplementary Fig. S6F**). The interactions between chain C of the spike and chain D of ACE2 (**Supplementary Fig. S6G**) displayed distinct patterns across different variants. WT and Beta followed similar interaction trends, with energy values ranging from −300 to 0 kcal/mol. Delta_*G*_ exhibited a positive interaction value during the simulation, which dropped to −400 kcal/mol after 0.75 *µ*s. The chain C of the spike showed no significant interaction with the chains E and F of ACE2 in none of the WT and variants (**Supplementary Fig. S6H-I**). Considering the pairing between the chains of the two proteins, the results are similar to those of the binding free energy, as illustrated through schematic representations in **Fig. 3G-J**. The most stable interactions happened along the direct interfaces (A-F, B-E and C-D), while in the variants less stable interaction values were observed with the neighboring chains. Specifically between chains C-F in Beta, B-D in Delta, and A-E and B-D in Delta_*G*_, considering a cut-off value of −50.0 kcal/mol. These results are in agreement with those of the binding free energy, suggesting more stable interactions between the spike and ACE2 in the variants compared to the WT.

### Conformational Rearrangement

We performed a cluster analysis to quantify the conformational rearrangement induced throughout the MD simulations. For this, we only considered the RBD and NTD domains of the spike in complex with the ACE2_*noT M*_ (see **Methods** section for the details of the clustering algorithm). Using a cutoff of 5 Å, we identified clusters for each studied systems and reported the population of the clusters (**Supplementary Fig. S7**). Considering highly populated clusters with cluster populations of at least 1000 frames, we extracted the representatives of each cluster. **Fig. 4A** presents the conformation of cluster representatives from each cluster for the WT, Beta, Delta, and Delta_*G*_ clusters. Each variant displays distinct conformational changes in the RBD and ACE2, critical for viral entry into host cells. One distinct observation relies on the number of clusters. While WT, Delta and Delta_*G*_ have 6, 7 and 5 clusters, respectively, Beta has the smallest number of clusters (only three), with one major cluster that includes 14,673 conformations out of 20,000 (containing 73.3 % of the trajectory). Moreover, analysis of the clusters sheds light on the rearrangements between different chains of spike and ACE2. In the case of WT, it is clear that the one-to-one interaction between chains A-F, B-E, and C-D remains stable along the simulations. However, additional chain interactions were observed for the variants. In the case of Beta, the RBD chain B is close to ACE2 chains D and E. At the same time, we observed very close distances between the RBD chain C and chains D and F of ACE2. This means the complex has undergone through conformational changes bringing RBD chain B closer to ACE2 chain D and RBD chain C closet to ACE2 chain F, while increasing the distances between RBD chain A and ACE2 chain E. In the case of Deta on the other hand, we observed a general rearrangement at the interface, which brings all three RBDs close to the three ACE2 proteins, in a way that each RBD is positioned close to its both ACE2 neighbors on each side, RBD chain A that is between ACE2 chains E and F, RBD chain B between both ACE2 chains D and E, and finally the RBD chain C at the interface of both ACE2 chains D and F. As for Delta_*G*_, a similar patter of forming a tight complex is observed at the interface of RBD chain A and ACE2 chain E and also RBD chain B and ACE2 chain D. However, the distance between RBD chain C and ACE2 chain F remains similar to that of the WT. These results are in agreement with binding free energy analysis findings and highlight better the conformational dynamics of the interaction formed between spike with three RBDs in upstate and in complex with three ACE2. The population of major clusters is reported in **Fig. 4B** for WT, Beta, Delta and Delta_*G*_.

**Figure 4:**
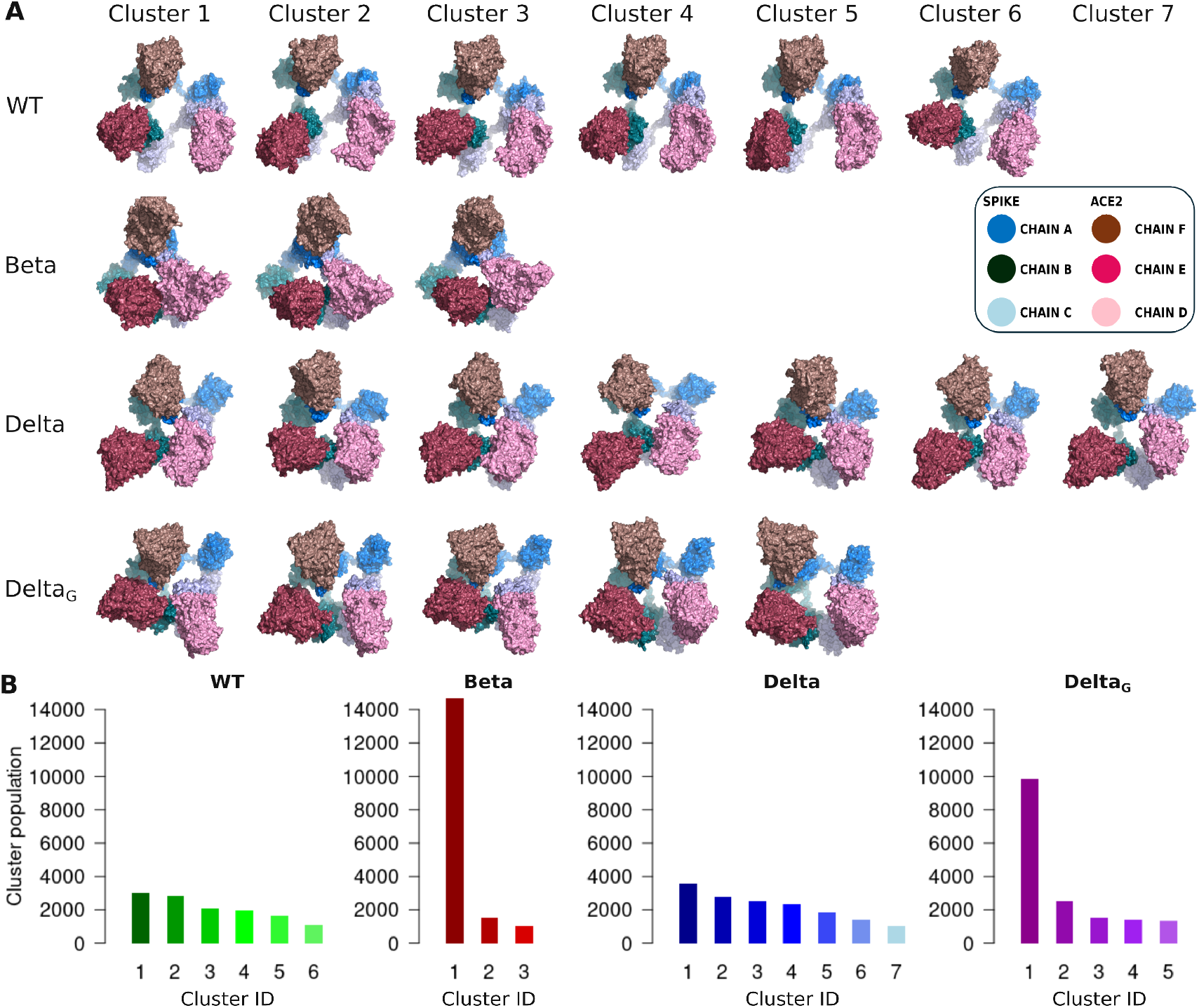
Conformational representatives for each studied system. **A)** The structure of representative conformations for the highly populated clusters is shown for WT, Beta, Delta, Delta_*G*_. Different shades of pink represent copies of ACE2, while shades of blue represent spike chains. **B)** The total number of conformations within the highly populated clusters are reported for each studied system.

### Role of Glycans

Glycans are carbohydrate molecules attached to spike proteins and play crucial roles in the function of the SARS-CoV-2 virus. Glycans contribute to protein folding and stability, forming a glycan shield that helps the virus evade immune detection. Considering the size of the system, and due to additional complexity and simulation time by adding glycans for all variants, we performed MD simulations in the absence of glycans for the WT, Beta and Delta variants, and investigated the role of glycans on stoichiometry only in the Delta_*G*_ variant. Differences between the Delta and Delta_*G*_ variants were observed in the H-bond formations. The Delta_*G*_ variant formed a unique H-bond, S477-M82 and E406-K353, absent in the Delta variant. Comparison of the salt bridges of the spike protein with the ACE2 interface of the Delta and Delta_*G*_ complexes showed that the Delta_*G*_ formed an additional salt bridge of E406-K353 and an additional interprotein interaction (K484-D216) between chains B and D of the spike protein and ACE2. The presence of glycans in Delta_*G*_ indicated that it formed another interaction network between the spike interface and ACE2. This analysis showed the critical role of glycans in the H-bond formations leading to stronger binding free energy and pair interaction, while consistent with the stoichiometry (1 RBD interaction with 2 ACE2) reported for Delta variant without any glycans.

## Discussion

The present study investigated the binding and conformational rearrangements between the spike trimer in a complex with three copies of ACE2. All the RBDs were in the open conformation, making interactions with their corresponding ACE2. We then performed MD simulations of the WT, Beta, Delta, and Delta_*G*_ variants, and investigated the generated trajectories to reveal the conformational dynamics of the complex. Our investigation into the H-bond network between the spike and ACE2 is of significant importance. The conservation or alteration of these H-bonds can profoundly impact the virus’s infectivity and immune escape capabilities.

In our study, WT formed the following H-bonds with the ACE2, Q493-K31, Q493-E35, T500-D355, G502-K353, D405-K353, D405-R559, R408-E564, and K417-D30, some of these interactions are similar to some of the notable H-bonds reported by [21]; their finding shows that several H-bonds were present at the WT RBD-ACE2 interface between residues Q493-E35, G502-K353, and K417-D30. WT formed an H-bond of K417-D30 between chain C-D, while the other variants have an observable mutation of K417N. [22] reported that T500-Y41, Q493-E35, and T500-D355 are the H-bonds formed between RBD and ACE2; these corroborate the H-bonds found in the Delta variant in our study. The Delta variant’s L452R, T478K, E484K, and N501Y mutations were not predicted to form H-Bonds at the interface. Delta_*G*_ formed additional H-bonds E406-K353, Y421-D30, G476-Q24, and S477-M82. The presence of these bonds may confer changes in the virus’s transmissibility within the host cell; each H-bond formed in Delta_*G*_ can potentially contribute to the overall affinity and stability of Delta_*G*_’s interaction with ACE2. Furthermore, residue K417 of WT formed a salt bridge with D30 of ACE2. This interaction was absent in the corresponding residue of the remaining variants due to the mutation K417N. [23] reported that K417 of SARS-CoV-2 formed a very stable salt bridge with D30 of ACE2. K417 provides a positively charged patch on the RBD of SARS-CoV-2. Similar to our study, [21] also reported the formation of salt bridges between the RBD residue K417 and the ACE2 residue D30 in WT.

Evaluating the binding energies of the Spike protein with the ACE2 for WT, Beta, Delta and Delta_*G*_ indicated that WT had a stronger binding affinity (−3.66 *±* 2.08 kcal/mol, **Supplementary Table S7**) for the ACE2 receptor between chains A-F compared to the other variant. Delta_*G*_ had a stronger binding affinity (−6.47 *±* 1.94 kcal/mol, **Supplementary Table S7**) for the ACE2 receptor between chains B-E than the other variant. In contrast, Beta had the strongest binding affinity (−5.11 *±* 1.88 kcal/mol, **Supplementary Table S7**) for the ACE2 receptor between chains C-D. The results also indicated that in addition to the initial interaction network between chains A-F, B-E, and C-D (one from spike and one from ACE2), other inter-protein interactions also happen along the simulations in Beta, Delta and Delta_*G*_.

Furthermore, the pair interactions showed that in addition to the initial interaction network between chains A-F, B-E, and C-D (one from spike and one from ACE2), other inter-protein interactions also happen along the simulations. Positive energy values indicate repulsive electrostatic interactions where charges repel each other, contributing positively to the system’s overall energy in the pair interaction calculation. The inter-protein interactions were observed mainly in the cases of Beta, Delta, and Delta_*G*_. Delta_*G*_ had a positive value for the pair interaction between chains C-D of the spike in complex with the ACE2 during the simulation and dropped to a negative value after 0.75 *µ*s; this implied that Delta_*G*_ moved from repulsive electrostatic interactions to attractive forces, indicating a more stable interaction between the Spike protein and ACE2. The pair interaction indicated the possibility of stoichiometry changes between the Spike and ACE2 during the simulations based on the inter-chain interactions observed in Beta, Delta and Delta_*G*_.

## Data availability

All the MD trajectories and final results of the case studied have been deposited in the Zenodo database (accession doi:10.5281/zenodo.14917210). We have also provided a Jupyter Notebook to reproduce the results of all the analysis reported in this study.

## Acknowledgments

This work was granted access to the HPC resources of IDRIS and CINES under the allocations 2021-AD010713015, 2022-A0130713799, 2023-AD010713015R1 and 2024-AD010713799R1 (granted to B.M, L.C. and Y.K.) made by GENCI. HK was supported by the French Agence Nationale de la Recherche (ANR), under grant ANR-22-CPJ2-0075-01.

## Notes

### Competing Interest Statement

The authors have declared no competing interest.

