## Supplementary Information for "Stoichiometric Insights into SARS-CoV-2 Spike–ACE2 Binding Across Variants"

### Supplementary Tables

Table S1: **RMSD values of individual chains.** The average and standard deviation values are reported for every chain of each system. Chains A, B and C are those of spike and chains D, E and F belong to the ACE2.

| Variants/RMSD (Å) | Chain A | Chain B | Chain C | Chain D | Chain E | Chain F |
|---|---|---|---|---|---|---|
| WT | 7.33 ± 0.96 | 7.68 ± 0.96 | 8.86 ± 1.36 | 6.45 ± 1.16 | 6.66 ± 0.88 | 7.25 ± 1.09 |
| Beta | 14.08 ± 1.79 | 7.27 ±1.09 | 6.24 ±0.94 | 7.40 ± 1.07 | 6.80 ± 1.27 | 8.03 ± 1.01 |
| Delta | 8.16 ± 1.08 | 9.11 ± 1.61 | 8.06± 1.05 | 5.60 ± 0.78 | 7.88 ± 1.40 | 6.57 ± 0.93 |
| $Delta_G$ | 6.77 ± 1.05 | 5.61 ±0.78 | 8.82 ±1.78 | 7.72 ± 1.72 | 6.17 ± 1.11 | 6.45 ± 1.38 |

Table S2: **RMSD values of the RBD domain and the $ACE2_{noTM}$.** The average RMSD and stand deviations are reported for both domains in each system. Chains A, B and C are those of spike and chains D, E and F belong to the ACE2.

| Variants/RMSD (Å) | Chain A | Chain B | Chain C | Chain D | Chain E | Chain F |
|---|---|---|---|---|---|---|
| WT | 3.63 ± 0.45 | 5.08 ± 0.62 | 5.46 ± 1.16 | 2.67 ± 0.27 | 3.08 ± 0.67 | 3.98 ± 0.56 |
| Beta | 4.75 ± 0.47 | 3.90 ± 0.43 | 3.53 ± 0.37 | 3.25 ± 0.31 | 3.49 ± 0.52 | 2.92 ± 0.34 |
| Delta | 4.28 ± 1.16 | 4.39 ± 0.92 | 3.35 ± 0.59 | 3.46 ± 0.43 | 3.72 ± 0.68 | 3.05 ± 0.29 |
| $Delta_G$ | 4.18 ± 0.61 | 3.95 ± 0.42 | 4.17 ± 0.75 | 3.37 ± 0.43 | 2.86 ± 0.28 | 2.45 ± 0.34 |

Table S3: **RMSF values of individual chains.** The average and standard deviation values are reported for every Chains of each system. Chains A, B and C are those of spike and chains D, E and F belong to the ACE2.

| Variants/RMSF (Å) | Chain A | Chain B | Chain C | Chain D | Chain E | Chain F |
|---|---|---|---|---|---|---|
| WT | 3.18 ± 1.70 | 2.99 ± 1.80 | 3.28 ± 1.81 | 2.27 ± 1.39 | 2.35 ± 1.29 | 2.14 ± 1.24 |
| Beta | 2.93 ± 1.50 | 2.67 ± 1.19 | 2.39 ± 1.02 | 2.14 ± 1.22 | 2.84 ± 1.85 | 1.87 ± 1.07 |
| Delta | 3.17 ± 1.78 | 3.28 ± 1.66 | 2.96 ± 1.47 | 2.11 ± 1.19 | 2.77 ± 1.65 | 2.15 ± 1.30 |
| $Delta_G$ | 2.74 ± 1.22 | 2.43 ± 1.25 | 2.84 ± 1.46 | 3.56 ± 1.83 | 2.72 ± 1.31 | 2.97 ± 1.58 |

Table S4: **RMSF values of the RBD domain and the ACE2$_{noTM}$.** The average RMSF and stand deviations are reported for both domains in each system. Chains A, B and C are those of spike and chains D, E and F belong to the ACE2.

| Variants/RMSF (Å) | Chain A | Chain B | Chain C | Chain D | Chain E | Chain F |
|---|---|---|---|---|---|---|
| WT | 1.39 ± 0.70 | 2.61 ± 1.57 | 1.91 ± 0.98 | 1.44 ± 0.83 | 1.68 ± 0.89 | 1.32 ± 0.75 |
| Beta | 1.22 ± 0.56 | 1.30 ± 0.71 | 1.29 ± 0.53 | 1.40 ± 0.63 | 1.75 ± 1.04 | 1.30 ± 0.71 |
| Delta | 1.91 ± 0.95 | 2.16 ± 1.13 | 1.98 ± 1.13 | 1.47 ± 0.80 | 1.91 ± 1.22 | 1,46 ± 0.78 |
| Delta$_G$ | 1.28 ± 0.64 | 1.49 ± 1.02 | 1.39 ± 0.74 | 1.47 ± 0.83 | 1.25 ± 0.67 | 1.24 ± 0.83 |

Table S5: **Key residues forming H-bonds.** The H-bonds formed at the interface of the Spike and ACE2 are reported from the WT, Beta, Delta, and Delta$_G$ simulations. Chains A, B and C are those of spike and chains D, E and F belong to the ACE2.

| Variants | Chain A-F | Chain B-E | Chain C-D |
|---|---|---|---|
| WT | Q493-K31, Q493-E35,T500-D355,G502-K353 | D405-K353 | D405-R559, R408-E564, K417-D30 |
| Beta | T500-Y41, G502-K353 | R403-D38, D405-K353, N487-Y83, G502-Q42 | R403-D38, Y453-D38, L455-HSD34, Y489-Y83, Q493-E35, T500-N64 |
| Delta | F486-Y83, Q493-K31, Q493-E35, T500-Y41, T500-D355, G502-K353 | R403-D38, D405-K353, G485-Y83, Y489-Y83, Q493-K31, Y505-D38 | T500-Y41, T500-D355, G502-K353 |
| Delta$_G$ | G502-K353 | R403-HSD34, R403-D38, D405-K353, E406-K353, Y421-D30, G476-Q24, S477-M82, Q493-K31 | R403-D38, Q493-K31, Q493-E35 |

Table S6: **Key residues forming salt bridges.** The salt bridges formed at the interface of the Spike and ACE2 are reported from the WT, Beta, Delta, and Delta$_G$ simulations. Chains A, B and C are those of spike and chains D, E and F belong to the ACE2.

| Variants | Chain A-F | Chain B-E | Chain C-D | Chain B-D |
|---|---|---|---|---|
| WT | K417-D30 | D405-K353, K417-E35 | K417-D30, D405-R559, D420-K26, R408-E564 | |
| Beta | | R403-D38, D405-K353 | R403-D38 | |
| Delta | | R403-D38, D405-K353 | | |
| Delta$_G$ | | R403-D38, D405-K353, E406-K353 | R403-D38 | K484-D216 |

Table S7: **Binding Free energy values.** The average and standard deviation values of binding free energy over the MD simulations are measured between the Spike and ACE2 chains of each system. Chains A, B and C are those of spike and chains D, E and F belong to the ACE2.

| Variants/Energy (kcal/mol) | WT | Beta | Delta | $Delta_G$ |
|---|---|---|---|---|
| Chain A-D | 0.00 | 0.00 | -0.34 ± 0.66 | 0.00 |
| Chain A-E | 0.00 | 0.00 | 0.00 | -0.80 ± 1.03 |
| Chain A-F | -3.66 ± 2.08 | -1.90 ± 1.40 | -2.87 ± 1.58 | -2.52 ± 1.69 |
| Chain B-D | 0.00 | -0.46 ± 0.57 | -1.55 ± 1.46 | -1.10 ± 1.14 |
| Chain B-E | -3.71 ± 1.60 | -3.86 ± 1.58 | -5.61 ± 1.86 | -6.47 ± 1.94 |
| Chain B-F | 0.00 | 0.00 | 0.00 | 0.00 |
| Chain C-D | -3.62 ± 1.56 | -5.11 ± 1.88 | -2.62 ± 1.90 | -4.89 ± 2.02 |
| Chain C-E | 0.00 | 0.00 | 0.00 | 0.00 |
| Chain C-F | 0.00 | -0.63 ± 0.91 | -0.21 ± 0.52 | -0.04 ± 0.19 |

Table S8: **Pair interaction values of the studied systems.** The average pair interaction values over the MD simulations are reported between the Spike and ACE2 chains of the studied systems. Chains A, B and C are those of spike and chains D, E and F belong to the ACE2.

| Variants/Pair interaction (kcal/mol) | WT | Beta | Delta | $Delta_G$ |
|---|---|---|---|---|
| Chain A-D | 0.00 | -0.16 ± 2.88 | -35.08 ± 51.91 | 0.00 |
| Chain A-E | -0.02 ± 0.23 | -0.03 ± 0.38 | -0.50 ± 4.42 | -55.38 ± 53.10 |
| Chain A-F | -195.60 ± 46.70 | -147.81 ± 48.45 | -142.89 ± 40.01 | -144.70 ± 34.32 |
| Chain B-D | -0.33 ± 5.15 | -29.17 ± 16.41 | -160.31 ± 120.34 | -87.31 ± 88.93 |
| Chain B-E | -222.86 ± 73.52 | -211.02 ± 47.54 | -205.26 ± 38.79 | -310.86 ± 64.81 |
| Chain B-F | 0.00 | 0.00 | 0.00 | 0.00 |
| Chain C-D | -270.13 ± 90.39 | -207.09 ± 48.51 | -179.02 ± 105.47 | -80.38 ± 290.04 |
| Chain C-E | 0.00 | -0.12 ± 0.41 | 0.00 | 0.00 |
| Chain C-F | 0.00 | -52.87 ± 32.14 | -22.46 ± 42.97 | -2.99 ± 14.88 |

### Supplementary Figures

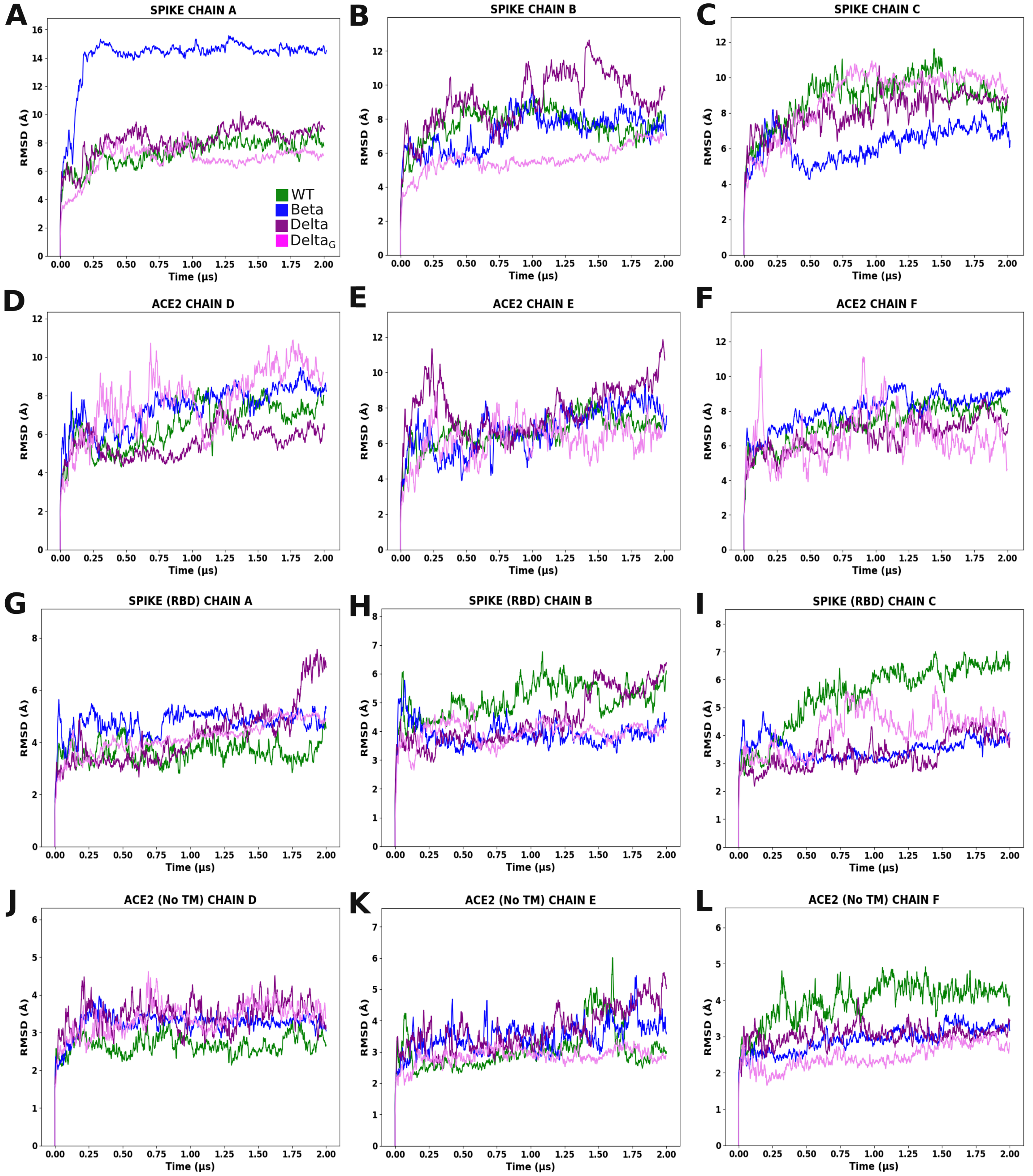


Figure S1: **RMSD Analysis of SARS-CoV-2 Variants. A-F)** displays the RMSD of individual chains, **G-I)** illustrate the RMSD of the RBD domain, and **J-L)** depict the RMSD of ACE2$_{noTM}$ region. The RMSD values are measured over the C$\alpha$ atoms with respect to the initial frame.

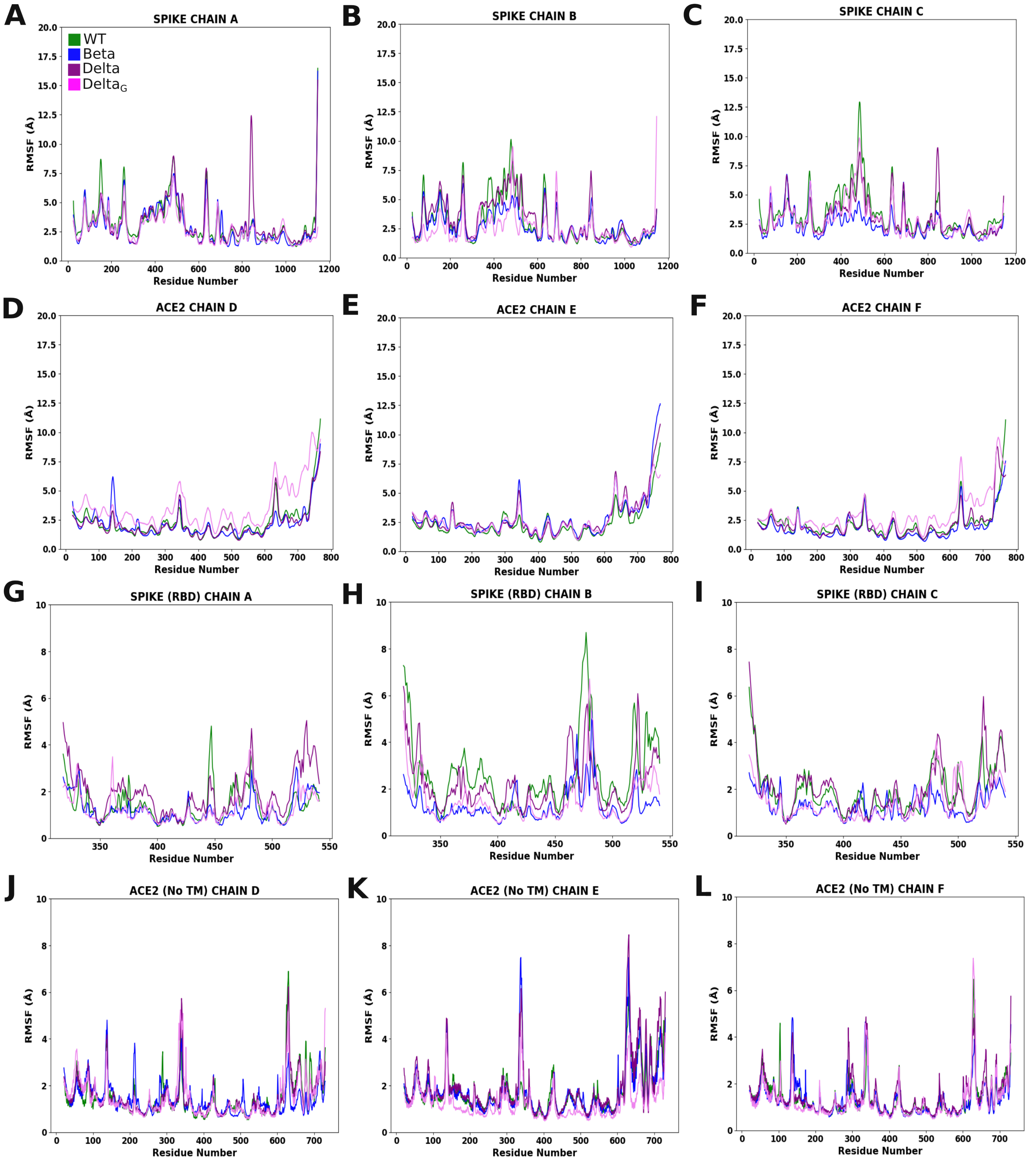


Figure S2: **RMSF Analysis of SARS-CoV-2 Variants. A-F)** displays the RMSF of individual chains, **G-I)** illustrate the RMSF of the RBD domain, and **J-L)** depict the RMSF of ACE2$_{noTM}$ region. The RMSF values are measured over the C$\alpha$ atoms and and with respect to the average confirmation.

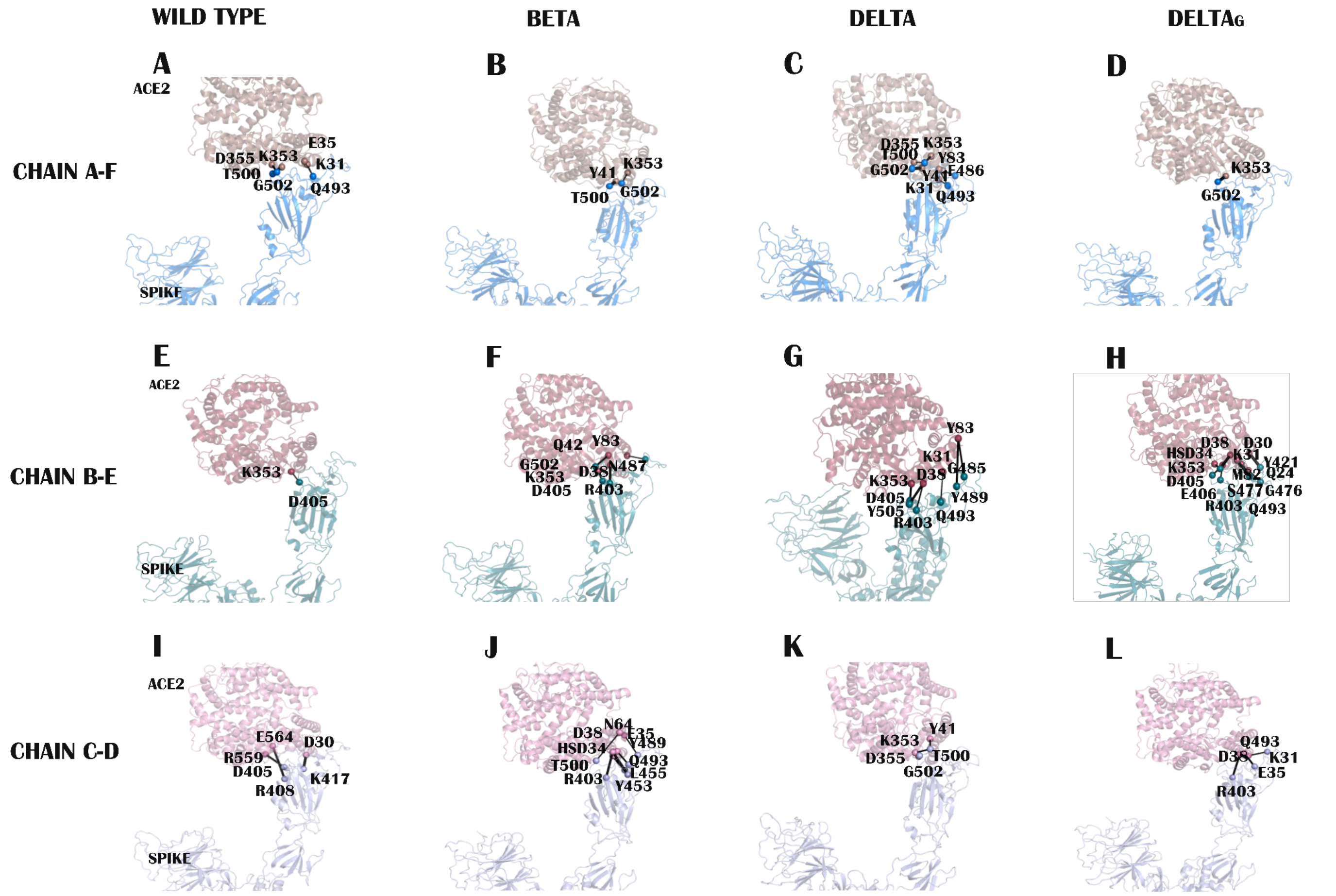


Figure S3: **Hydrogen Analysis of SARS-CoV-2 Variants.** The key residues involved in the H-bonds formed at the interface between the chains of spike and ACE2 are shown for the WT (**A, E, I**), Beta (**B, F, J**), Delta (**C, G, K**), and Delta$_G$ (**D, H, L**).

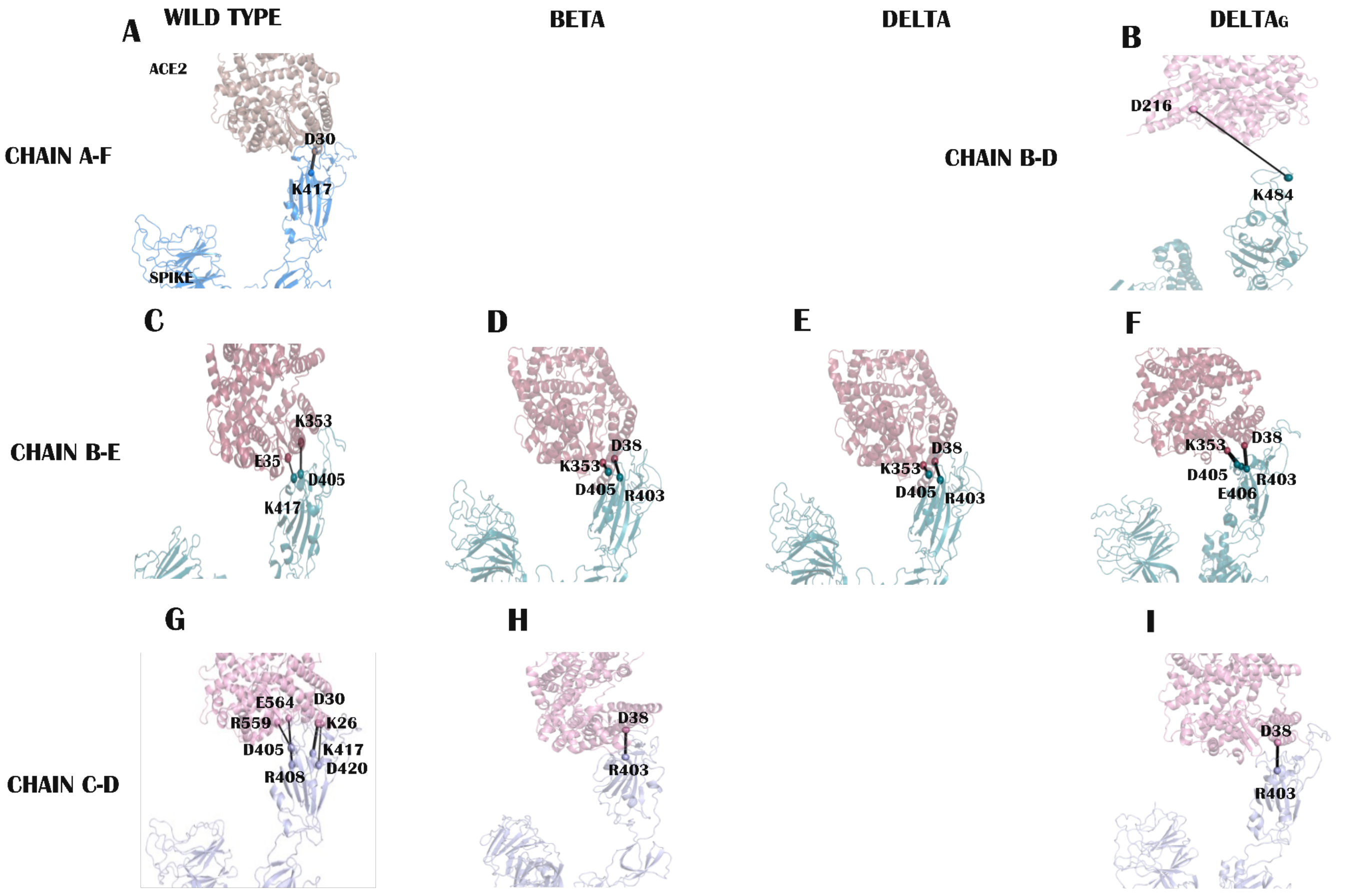


Figure S4: **Salt bridge Analysis of SARS-CoV-2 Variants.** The key residues involved in the H-bonds formed at the interface between the chains of spike and ACE2 are shown for the WT (**A, C, G**), Beta (**D, H**), Delta (**E**), and Delta$_G$ (**B, F, I**).

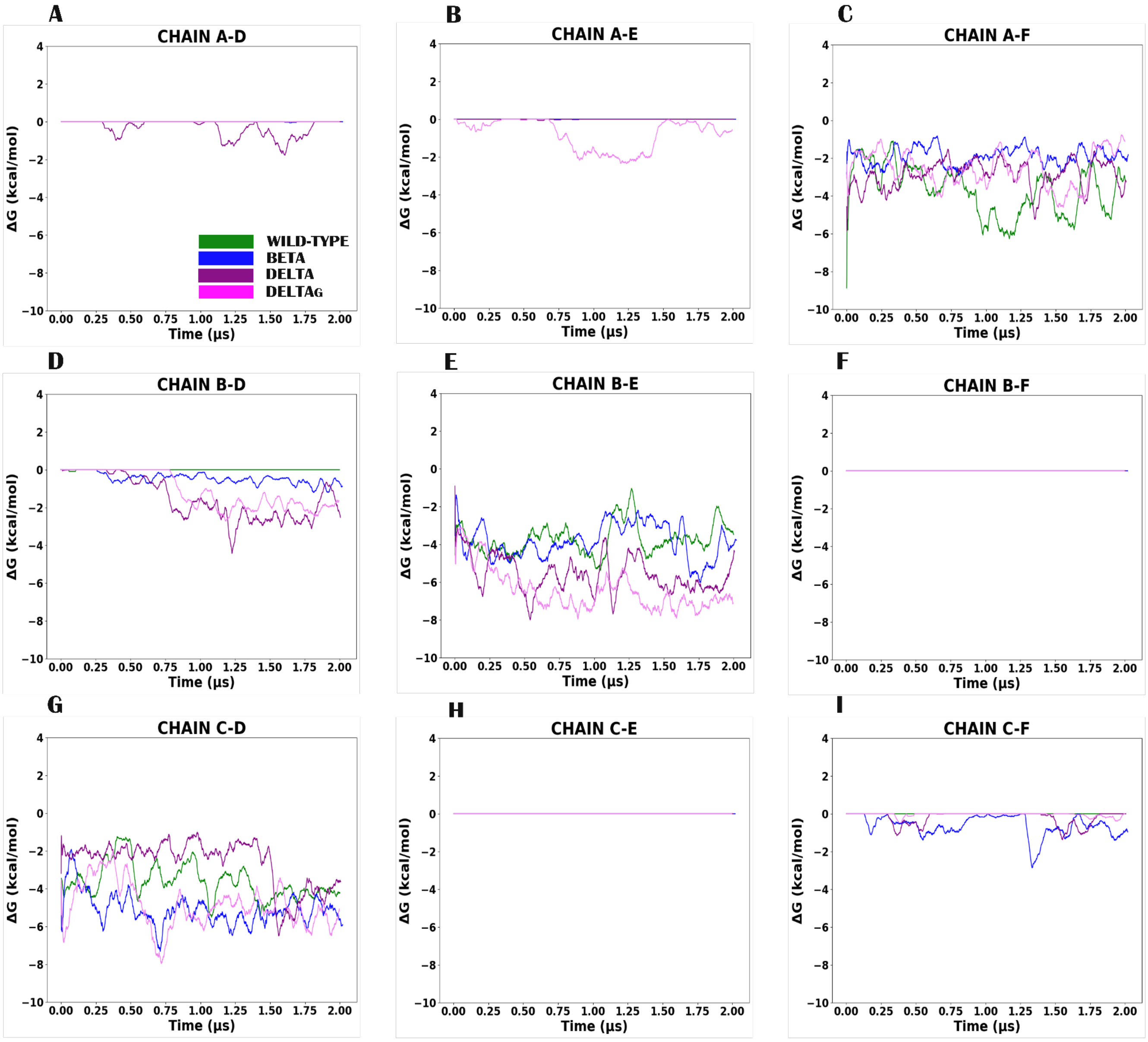


Figure S5: **Binding Free Energy Analysis.** The changes of binding free energy between the chains of spike and ACE2 are reported over the course of the MD simulations, for the WT (green), Beta (blue), Delta (purple) and Delta$_G$ (pink) variants. Chains A, B and C are those of spike and chains D, E and F belong to the ACE2. The initial one-to-one interactions between spike and ACE2 was formed between **C)** chains A-F, **E)** chains B-E and **G)** chains C-D. The other subplots represent binding free energy between chains that were not directly in contact.

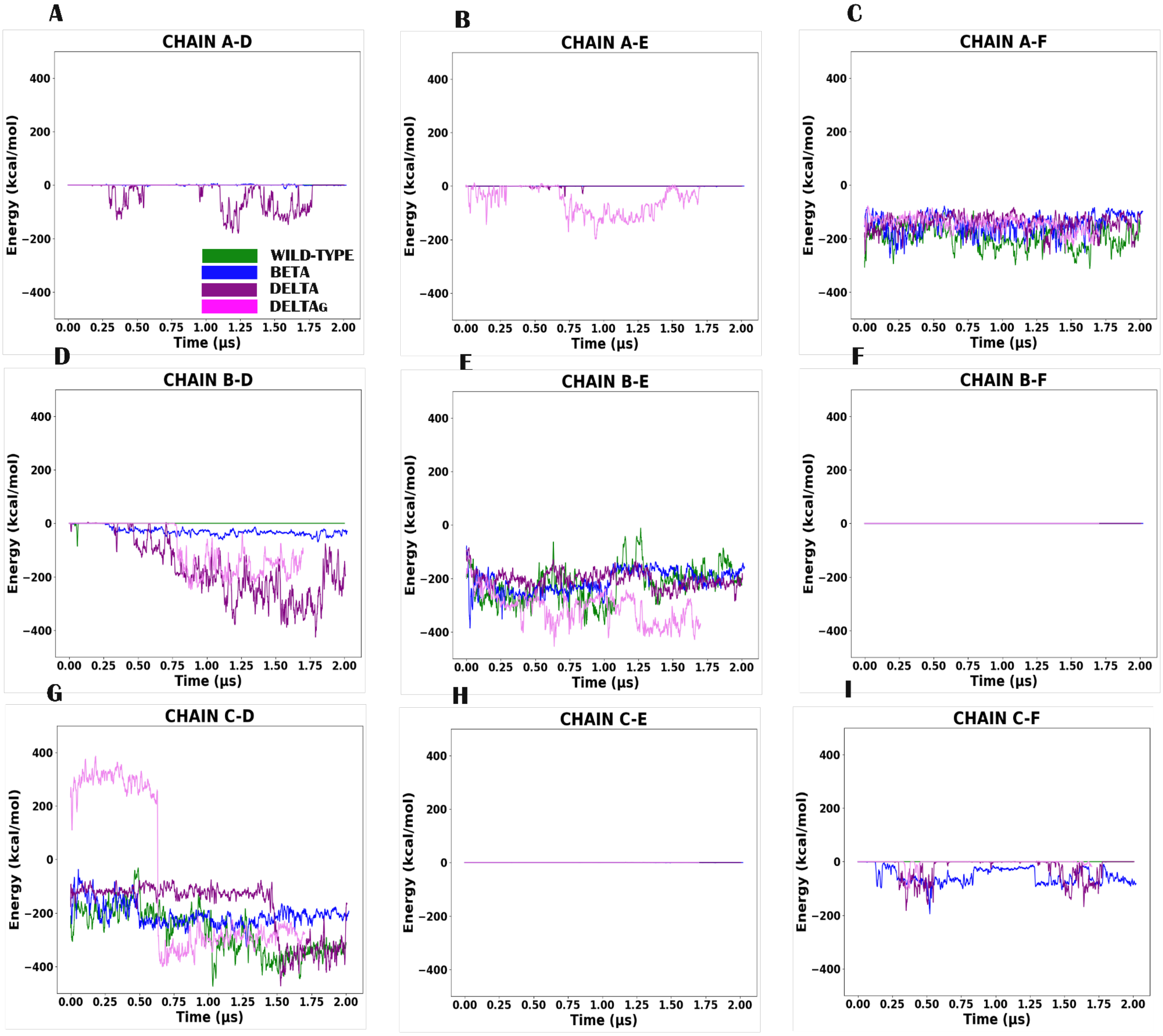


Figure S6: **Analysis of pair interactions.** Pair interaction values between the chains of spike and ACE2 are reported for the WT (green), Beta (blue), Delta (purple) and Delta$_G$ (pink) variants. Chains A, B and C are those of spike and chains D, E and F belong to the ACE2. The initial one-to-one interactions between spike and ACE2 was formed between **C)** chains A-F, **E)** chains B-E and **G)** chains C-D. The other subplots represent pair interactions between chains that were not directly in contact.

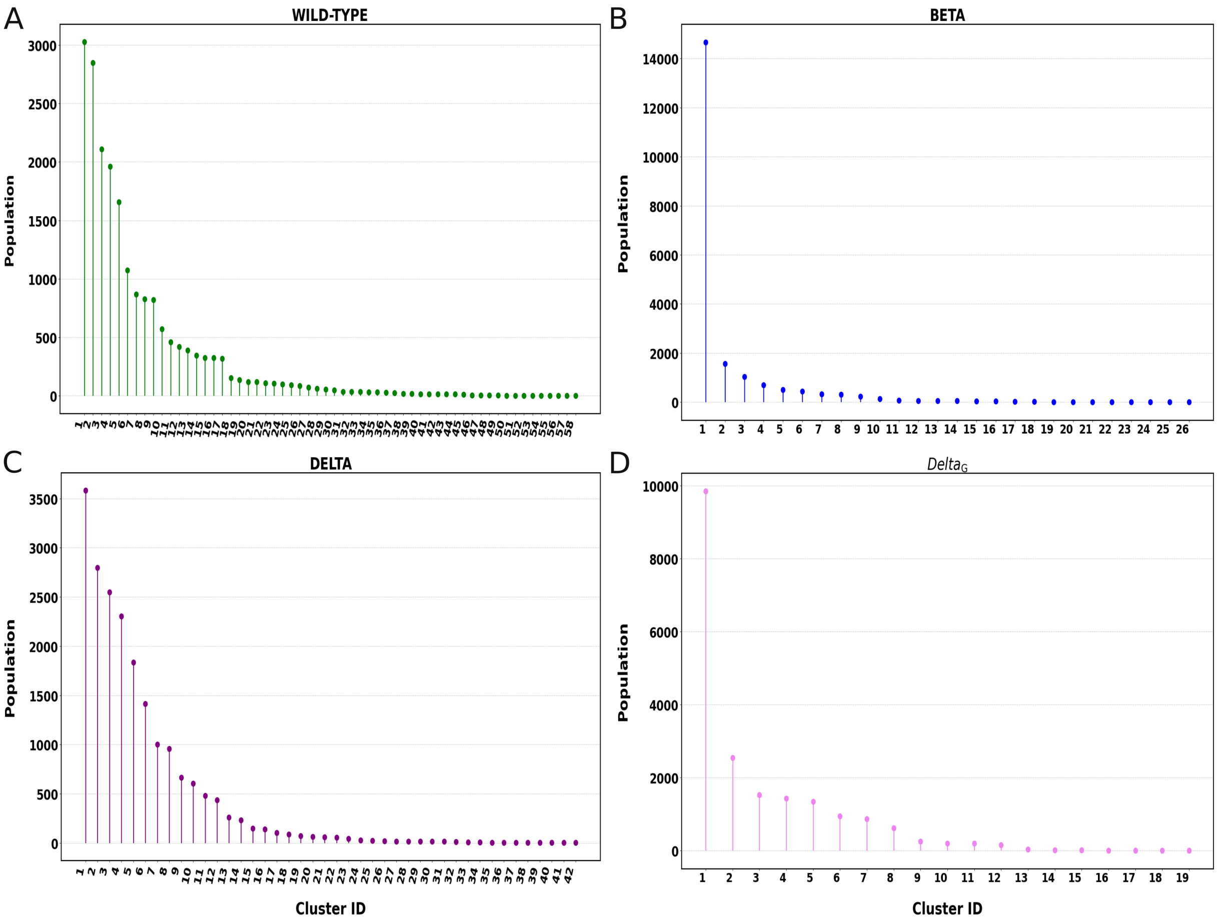


Figure S7: **Details of the cluster analysis.** The cluster populations are reported for **A)** WT, **B)** Beta, **C)** Delta, and **D)** $\mathrm{Delta}_G$.
